# Gene symbol recognition with *GeneOCR*

**DOI:** 10.1101/2022.07.01.498459

**Authors:** Paul Geser, Jitao David Zhang

## Abstract

**Motivation:** Figures in biomedical publications often contain lists of gene symbols. Human intelligence and manual curation is required to recognize, extract, and validate the symbols in order to reuse them. The task is laborious, error prone, and amenable to automation.

**Results:** We introduce *GeneOCR*, a web-based software to recognize, extract, and check gene symbols from figures. It features both an interactive user interface and programmatic access. Simulation-based benchmark reveals that *GeneOCR* recognizes or suggests correct gene symbols in more than 80% cases. The performance qualifies *GeneOCR* as an enhancement to manual curation.

**Availability:** The source code of *GeneOCR* is available with GNU v3 license at https://github.com/bedapub/GeneOCR-UI (frontend) and https://github.com/bedapub/GeneOCR-API (backend). A Docker/Singularity image is available via *BioContainers*.

## 1 Introduction

Biomedical publications such as journal papers, presentations, and posters often contain figures showing a list of gene symbols. Researchers often need to extract lists of gene symbols from publications manually and store them in a machine-readable format such as plain text file in order to re-use the list for their own work. In few cases, for instance with a vector-based PDF file, it is possible to extract the gene symbols simply with copy/paste. In most other cases, however, when the figure is only available in a raster-based PDF file or pixel formats such as PNG, JPEG and TIFF, human intelligence and manual work is required to recognize and extract the gene symbols. In either case, additional work is often necessary to check the extracted symbols as valid identifiers. The laborious and error-prone process does not scale up with the increasing need of knowledge accumulation and reuse. While tools like *Google Keep* are able to extract texts from images, to our best knowledge there is no open-source tool to automate the task of gene symbol recognition and validation.

To close the gap, we introduce the software *GeneOCR* (OCR=optical character recognition). It employs a state-of-the-art character recognition system to recognize gene symbols from figures. *GeneOCR* offers both a front-end for end users and application programming interfaces (APIs) to allow programmatic access.

### 2 Software features

*GeneOCR* offers the following functionalities:

1. Optical character recognition (OCR): *GeneOCR* recognizes and extracts texts in images. User can select gene symbols or free text as input.
2. Enhancement utilities for OCR:
  a. Rotating: If the texts are not horizontally aligned, *GeneOCR* rotates the image automatically.
  b. Multiple cropping: User can analyze several regions of the same figure with *GeneOCR*.
  c. Optional sharpening: Blurry images may reduce the accuracy of OCR. *GeneOCR* sharpens the image upon request.
3. Spell check: if the input texts are gene symbols, *GeneOCR* checks the extracted symbols against with a dictionary of valid gene symbols of commonly used model species provided by NCBI Gene (Gene, 2022). *GeneOCR* highlights invalid gene symbols and suggests similar, valid symbols as alternatives.
4. Downloading: The user can download extracted gene symbols or free texts in plain text or other formats.

## 3 Software architecture

The *GeneOCR* software consists of two layers: the web-browser-based frontend, implemented with *ReactJS*, and the Python-based backend, implemented with *FastAPI*. The backend depends on version 5 of the open-source *Tesseract* engine (Tesseract, 2022), which uses long-short term memory (LSTM, Hochreiter and Schmidhuber, 1997), an artificial neural network, for text recognition. The backend also uses *OpenCV*, a real-time optimized computer vision library, for image processing. The simple yet robust two-layer model allows further extension and automation of tasks via API calling. *GeneOCR* is packaged and distributed in Docker/Singularity images that run virtually in any modern computer environment via *BioContainers* (da Veiga Leprevost *et al*., 2017).

## 4 Example

Figure 1 demonstrates the interactive use of *GeneOCR*. In step 1, the user uploads an image and manually crops one or more text areas that contain gene symbols (blue box in the figure). In step 2, *GeneOCR* performs text recognition, highlights invalid ones, and proposes similar symbols as alternatives. In step 3, the user can change recognized symbols either by accepting a suggestion, or by manual editing, in which case the validity is also checked. Once satisfied with the results, the user can download them in a file.

**Figure 1.**
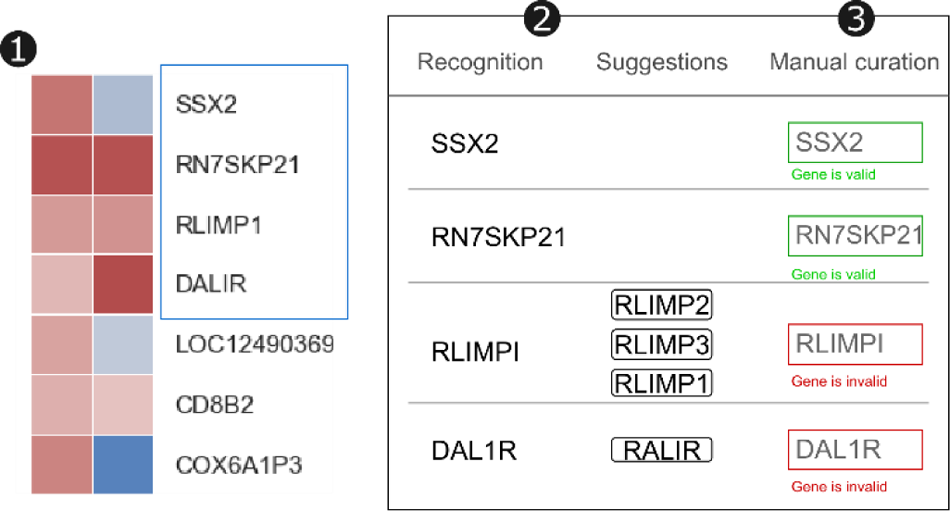
A case study of working with *GeneOCR*, showing the three steps described in the *Example* section.

## 5 Benchmark with simulation

We investigated the performance of *GeneOCR* with simulations. To this end, we created figures containing 307 randomly selected human and mouse gene symbols, ran *GeneOCR*, and compared the output with input. Benchmark suggests that *GeneOCR* has a satisfactory performance. It recognizes the correct symbols in 65% cases. Among symbols with recognition errors, *GeneOCR* suggests correct items in 57% cases. The errors affect mostly one character: the mean length-normalized Levenshtein distance between the output with error and the input is 0.21, with 95% confidence interval 0.14-0.27, *i*.*e*. one wrong character in a 4-7 character symbol. The errors are mostly due to substitution of optically similar characters, *e*.*g. 1* for *I* or *O* for *0*. In summary, *GeneOCR* recognizes or suggests the correct gene symbol in >80% cases and the errors in the rest case involve mostly single characters.

## 6 Conclusions

*GeneOCR* recognizes and extracts gene symbols in visual elements of scientific publications. With an intuitive user interface, programmatic access, and a decent performance, *GeneOCR* simplifies and enhances manual curation. We invite researchers to apply and co-develop *GeneOCR*, build tools and pipelines using it, and send us feedback.

## Acknowledgements

We thank Fabian Birzele, Hans Friedrich, Ralf Horstmöller, and Matthias Nettekoven for the support of the apprenticeship of Paul Geser. We thank colleagues of the Predictive Modeling and Data Analytics chapter for feedback.

